# Robotic Visible-Light Optical Coherence Tomography Visualizes Segmental Schlemm’s Canal Anatomy and Segmental Pilocarpine Response

**DOI:** 10.1101/2024.09.23.614542

**Authors:** Raymond Fang, Pengpeng Zhang, Daniel Kim, Junghun Kweon, Cheng Sun, Alex S. Huang, Hao F. Zhang

## Abstract

**Purpose:** To use robotic visible-light OCT (vis-OCT) to study circumferential segmental Schlemm’s canal (SC) anatomy in mice after topical pilocarpine administration.

**Methods:** Anterior segment imaging was performed using a vis-OCT sample arm attached to a 6-degree-of-freedom robotic arm to maintain normal (perpendicular) laser illumination aimed at SC around the limbus. Sixteen mice were studied for repeatability testing and to study aqueous humor outflow (AHO) pathway response to topical drug. Pharmaceutical-grade pilocarpine (1%; n = 5) or control artificial tears (n = 9) were given, and vis-OCT imaging was performed before and 15 minutes after drug application. After SC segmentation, SC areas and volumes were measured circumferentially in control- and drug-treated eyes.

**Results:** Circumferential vis-OCT provided high-resolution imaging of the anterior segment and AHO pathways, including SC. Segmental SC anatomy was visualized with the average cross-sectional area greatest temporal (3971 ± 328 µm^2^) and the least nasal (2727 ± 218 µm^2^; p = 0.018). After pilocarpine administration, the iris became flatter, and SC became larger (pilocarpine: 26.8 ± 5.0% vs. control: 8.9 ± 4.6% volume increase; p = 0.030). However, the pilocarpine alteration was segmental as well, with a greater increase observed superior (pilocarpine: 31.6 ± 8.9% vs. control: 1.8 ± 5.7% volume increase; p = 0.023) and nasal (pilocarpine: 41.1 ± 15.3% vs. control: 13.9 ± 4.5% volume increase; p = 0.045).

**Conclusion:** High-resolution circumferential non-invasive imaging using AS-OCT of AHO pathways is possible in living animals with robotic control. Segmental SC anatomy was seen at baseline and was consistent with the known segmental nature of trabecular AHO. Segmental SC anatomical response to a muscarinic agonist was seen as well. Segmental glaucoma drug response around the circumference of AHO pathways is a novel observation that may explain the variable patient response to glaucoma treatments.

## Introduction

Glaucoma is an optic neuropathy and a leading cause of irreversible blindness worldwide^1^. The only FDA-approved treatment is to lower intraocular pressure (IOP)^2–4^ because no existing glaucoma therapy directly treats the optic nerve. Instead, glaucoma is treated by risk factor modification, and elevated IOP is the only risk factor that can be modified either by drops, lasers, or surgeries^2–4^.

IOP is modeled by the Goldmann equation, where the final pressure arises from a balance between aqueous humor production and aqueous humor outflow (AHO)^5^. There are multiple AHO pathways, and the trabecular pathway is the primary source of AHO resistance (> 50%)^6^. In the trabecular pathways, aqueous flows from the anterior chamber (AC) to the trabecular meshwork (TM), through Schlemm’s canal (SC), and then into distal outflow pathways, including collector channels (CCs), the intracellular venous plexus, and aqueous/episcleral veins before returning aqueous to the blood circulation^7, 8^. However, this traditional linear description does not fully describe AHO circumferentially around the limbus in a three-dimensional (3D) eye.

Recent evidence by multiple groups has shown that circumferential trabecular AHO is segmental. In the laboratory, perfusion of fluorescent microbeads has demonstrated segmental AHO with TM bead trapping in high-flow (HF) and low-flow (LF) regions^9–13^. These TM regions then express different proteins and extracellular matrix (ECM) and display varying tissue stiffness^9–12^. Imaged on the ocular surface, aqueous angiography uses soluble tracers and has shown segmental post-limbal HF and LF regions in multiple species as well^14–22^. Since aqueous angiography can be performed in patients^23–27^, it holds promise in influencing glaucoma patient care. However, aqueous angiography is limited because it is laborious, time-consuming, and invasive, resulting in risks. Thus, a non-invasive imaging approach is preferred.

While OCT is a natural candidate to address the above need^28, 29^, OCT has mostly been used for the posterior segment. Currently, anterior-segment OCT (AS-OCT) is limited to imaging the large, complex, and circular AHO pathway anatomy. AHO pathways are deeply positioned (∼50-500 µm) within highly scattering sclera, making illumination penetration and imaging depth a limiting factor for commercial AS-OCTs. AHO pathway structures such as CCs can be small (∼10 microns) and below commercial AS-OCT resolution^30, 31^. Also, given the circular post-limbal structural organization of AHO pathways, the circumferential distance (>37 mm) and area (>200 mm^2^) to be imaged are beyond the field-of-view (FOV) of current AS- OCTs.

These anatomical features also prevent stationary imaging devices from maintaining normal (e.g., perpendicular) illumination along the entire circular AHO pathway organization. In OCT, normal illumination is essential for minimizing optical pathlength within the tissue and the resulting light scattering and maximizing the inherent high axial resolution of OCT to achieve accurate cross-sectional imaging. Some commercial AS-OCTs require re-positioning cameras while patients take eccentric gazes to achieve normal illumination^32^. However, eccentric gazes are challenging to hold, and moving both the eye and the camera sacrifices reproducibility during longitudinal study. Imaging a relaxed and forward-looking subject is better.

Despite all of these challenges, AS-OCT has been reported for AHO pathways. We previously conducted a 360-degree reconstruction of AHO pathways in the human eye using overlapping volumes acquired from a commercial AS-OCT^32^. However, in this study, imaging depth was limited, oblique illumination was used, and both AS-OCT and ocular re-positioning were necessary. Image acquisition took days as >5000 B-scans were required, and only one subject could be imaged. Therefore, current AS-OCTs cannot feasibly image the entire circumferential AHO pathways. This is consistent with reported AHO AS-OCT studies where imaging in small areas is extrapolated to study the entire circumferential anatomy, leading to a limited understanding of AHO^33–36^.

Therefore, to overcome challenges in imaging AHO pathways, we developed a robotic visible light-optical coherence tomography system (vis-OCT) and demonstrated its utility in imaging the full 360 degrees of the AHO pathways in an *in- vivo* mouse eye^37^. We addressed the attenuation of visible light by angling the incident light so that it traveled the shortest distance to reach AHO pathways. To optimize the visualization of the SC, we used robotics to position our device such that the OCT beam was parallel to the minor axis of the SC around the full circumference of the eye. Using this tool, we now evaluate the *in-vivo* anatomical morphology of SC in the living mouse eyes at baseline and after pharmacological stimulation. We hypothesize that the baseline AHO pathway anatomy is segmental and that the physiological response to pilocarpine^38, 39^, a drug known to decrease AHO pathway resistance and lower IOP, is also segmental.

## Methods

### Animal Preparation

All experimental protocols were approved by the Northwestern University Institutional Animal Care and Use Committee and complied with the ARVO Statement for the Use of Animals in Vision Research. Sixteen adult wild-type C57BL/6J mice were used for our experiments. Two mice were imaged for repeatability assessment. An additional 14 mice were imaged before and after drug treatment and compared. We kept the mice under a 12-hour light/12-hour dark cycle with unrestricted access to food and water in the Center for Comparative Medicine at Northwestern University.

Prior to imaging and any procedures, mice were anesthetized using intraperitoneal injection (10 mL/kg body weight) with a ketamine/xylazine cocktail (ketamine: 11.45 mg/mL; xylazine: 1.7 mg/mL, in saline). Body temperature was maintained using a heat lamp.

### 360 Degrees vis-OCT imaging and data reconstruction

We used a custom-developed robotic visible-light optical coherence tomography (vis-OCT) device to image the full 360 degrees of the AHO pathway (Fig. 1a)^40^. Our vis-OCT system operated between 510 nm and 610 nm^41^. The theoretical axial resolution of vis-OCT was 1.3 µm in tissue, and the theoretical lateral resolution was 9.4 µm^40^. To expose the entire AHO pathways for imaging, we made two relaxing incisions at the nasal and temporal canthi and inserted a circular speculum (Mouse Circular Speculum Type SS, Focus Ophthalmics) underneath the eyelid. Next, we acquired eight volumetric vis-OCT scans around the circumference of the mouse eye (Fig. 1b), where the robotic arm rotated the incident OCT beam 45 degrees around the eye’s optical axis between each acquisition. We set the lateral FOV for each OCT volume as 2.04 mm. Each volume consisted of 512 A-lines per B-scan and 512 B-scans per volume. We acquired each volume using a temporal speckle averaging data acquisition pattern^42^, where each B-scan was repeated twice per volume, and three volumes were acquired. We acquired the data at an A-line rate of 75 kHz and an incident beam power of 1mW. All quadrants of the mouse eye were at the same height during image acquisition.

**Figure 1.**
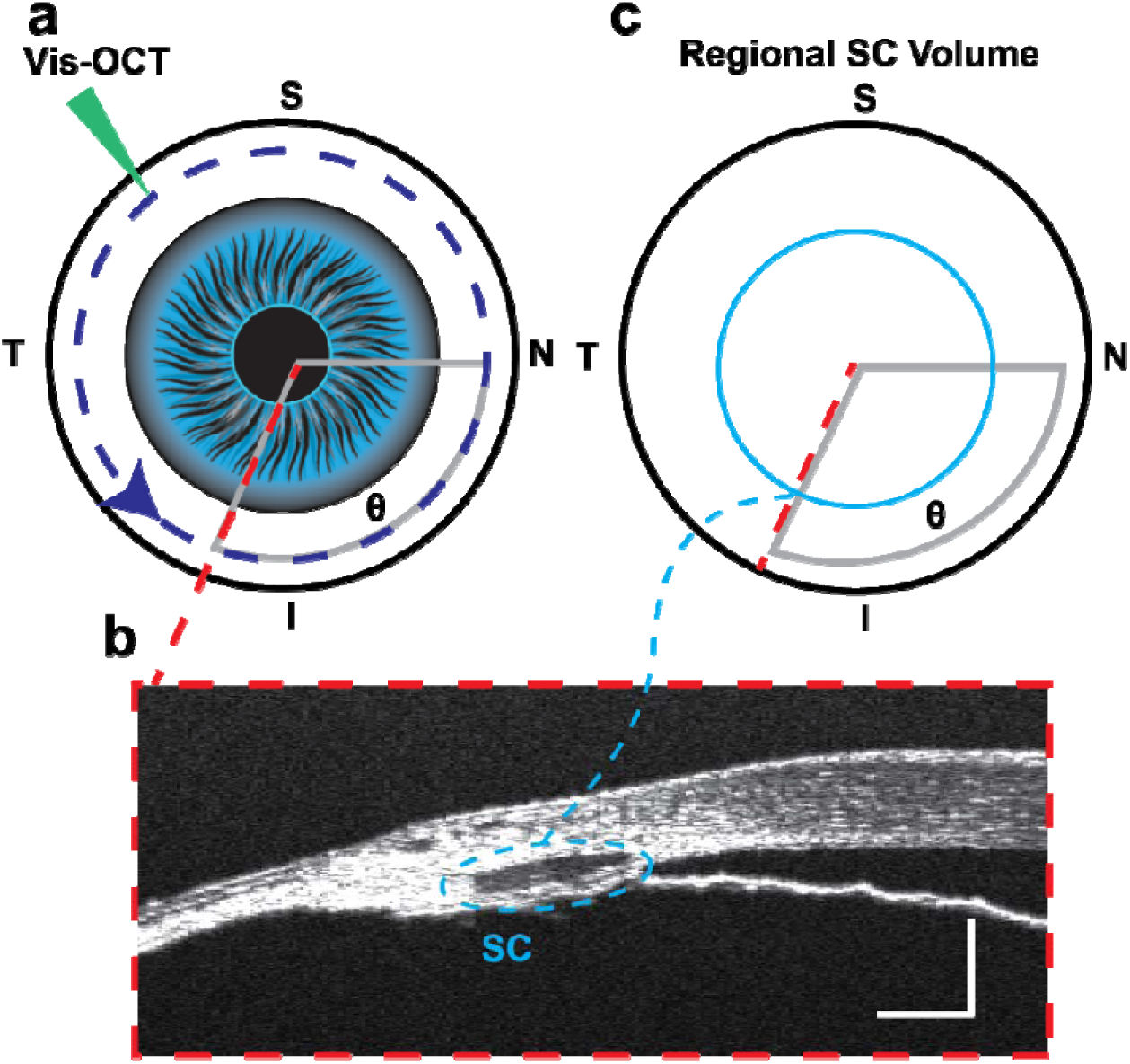
Vis-OCT characterizes SC morphology around the entire circumference of the globe. **(a)** Illustration of robotic vis-OCT imaging of the limbus of the eye. **(b)** A representative B-scan image of the limbus at an angle θ relative to the nasal (N) quadrant. The SC is the hypo-reflective lumen within the blue dotted oval. **(c)** The cross-sectional SC was assessed relative to the angular position (blue dotted line). Scale bars are 100 μm.

### Schlemm’s canal size measurement

We determined the orientation for each of the eight acquired vis-OCT volumes relative to the eye’s optical axis. Based on the relative angular orientation of each volume to the nasal canthus, where the upper and lower eyelid meet on the medial corner of the eye, we assigned an angular orientation of each cross-sectional B-scan relative to the nasal quadrant of the eye (angle θ; Fig. 1c). For each volume, we segmented SC for one out of every thirty cross-sectional B-scans acquired, amounting to a separation of 120 µm per segmented cross-section. We calculated the SC cross-sectional area per degree of the eye by averaging the cross-sectional areas of segmented B-scans located within twenty degrees of the plane corresponding to each angular position (angle θ ; Fig. 1c). Additionally, we calculated the SC volume by summing the cross-sectional areas of SC for every B-scan within the region of interest.

### Measuring SC morphology changes in response to pilocarpine

We imaged the entire outflow pathway at baseline. Fig. 2a shows an example of a digitally resampled AS-OCT scan for a C57BL/6 mouse before and after pilocarpine. As previously described^37^, we performed digital resampling by viewing a curved plane passing through the center of SC in the 3D processed data and flattening the outer surface of the resulting image. Briefly, we used the skeleton of the segmented SC to determine the trajectory of the curved plane. Following imaging of the entire anterior segment, we added a 1% pilocarpine eye drop (n = 5 eyes) (NDC 61314- 203-15, Sandoz). After 15 minutes, we reimaged the anterior segment again. Fig. 2b shows the same AS-OCT view from the same eye following pilocarpine administration. We calculated the SC cross-sectional area per degree of the eye at baseline and after drug application (Fig. 2c). For each angular degree, we calculated the percent size change, defined as the difference in SC cross-sectional area after eye drop administration divided by the initial cross-sectional area at baseline (Fig. 2d). We performed control experiments using the same methodology, except that artificial tears (n = 9) (NDC 57896-181-05, Gericare) were used.

**Figure 2.**
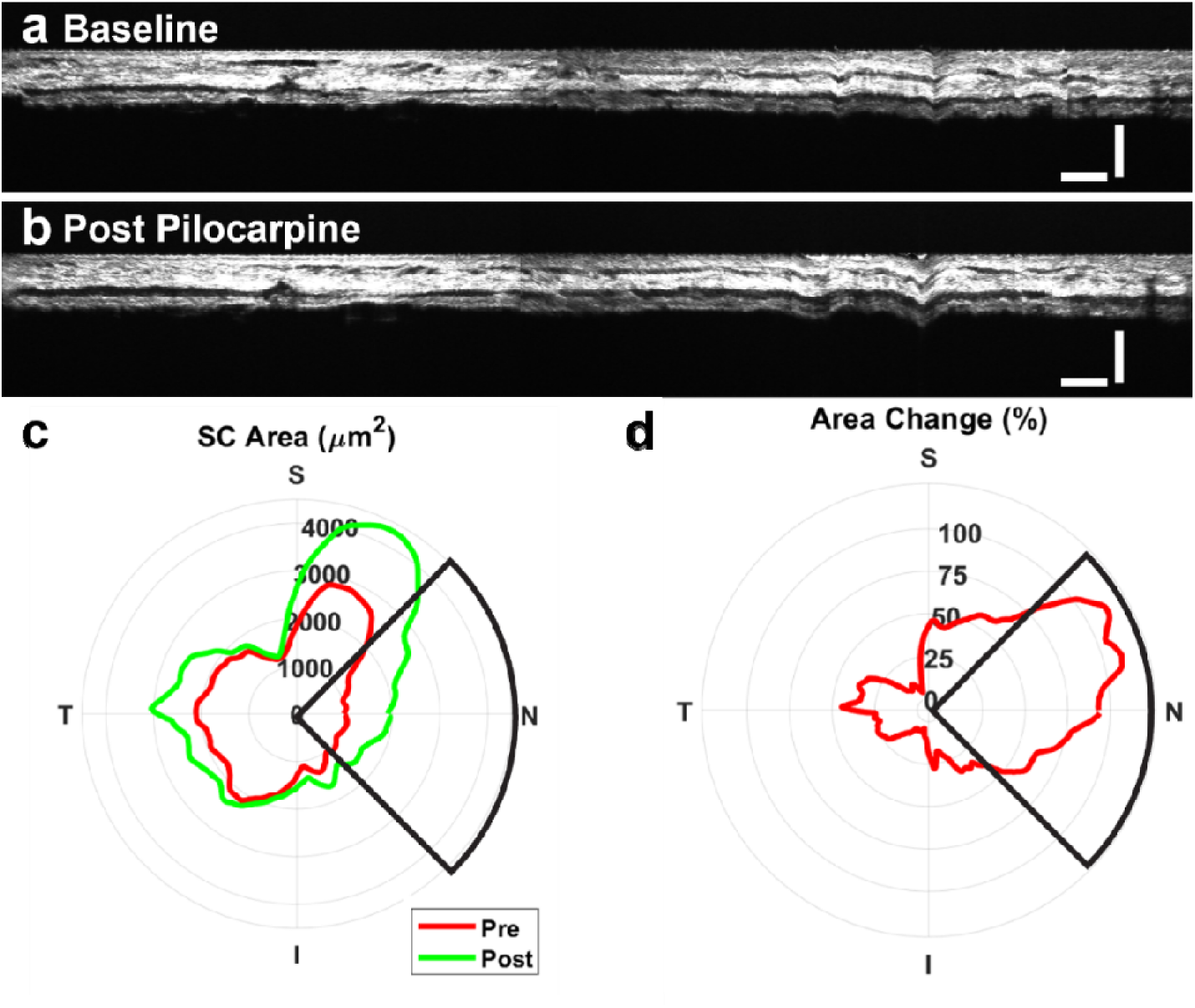
SC morphology changes segmentally in response to pilocarpine. **(a)** Digitally resampled B-scan image before pilocarpine administration. **(b)** Digitally resampled B-scan image 15 minutes after pilocarpine administration. **(c)** SC cross-sectional area per degree of eye before and after administration of pilocarpine. **(d)** Percent increase in SC area per angle of the eye relative to the initial volume at a given circumferential location. OCT in (a) and (b) correspond to the nasal pie wedges highlighted in (c) and (d). Scale bars are 100 μm.

### Statistical Analysis

We applied an unpaired t-test for all single comparisons, including the percent volume change and the average standard deviation of clock hour volume change per eye after applying 1% pilocarpine or artificial tears. To compare the segmental differences between the eye quadrants and clock hours, we ran a one-way ANOVA. We used the Sidak method to correct for multiple comparisons. Measurements are reported as mean±standard error (SEM) unless specified otherwise.

## Results

### Vis-OCT visualizes SC and demonstrates segmental morphology

To assess the repeatability SC measurements, we imaged the same eye region and quantified the average SC cross-sectional area for a single volume taken multiple times, 5 minutes apart, in two C57BL/6 mice. In the first mouse eye, the average difference between 4 repeated measures of SC cross-sectional area was 0.4±2.6%. In the second mouse eye, the average difference across 3 repeated measures of SC cross-sectional area was -0.5±3.5%.

Given that AHO is segmental, we first sought to test whether segmental patterns in SC anatomy existed. Fig 3a shows the average SC cross-sectional area for every eye quadrant, with the average SC cross-sectional area in the temporal quadrant being larger than in the nasal quadrant (p=0.018; n=14). We measured an average SC cross-sectional area of 2727±218 µm^2^ for the nasal quadrant, 3187±313 µm^2^ for the superior quadrant, 3971±328 µm^2^ for the temporal quadrant, and 3018±257 µm^2^ for the inferior quadrant. To account for individual baseline variation, Fig. 3b plots the average cross-sectional area for each quadrant divided by the average area across each entire eye, with the relative SC cross-sectional area largest in the temporal quadrant (p-values: <0.001-0.031; comparing the temporal quadrant to other quadrants). We found the SC cross-sectional area relative to the mean across all quadrants to be 0.85±0.06 for the nasal quadrant, 0.98±0.07 for the superior quadrant, 1.23±0.05 for the temporal quadrant, and 0.94±0.06 for the inferior quadrant. We note that the segmental pattern of SC area was different for each eye, with the relative SC area ranging from 0.64 to 1.42 for the nasal quadrant, 0.41 to 1.34 for the superior quadrant, 0.77 to 1.44 for the temporal quadrant, and 0.43 to 1.22 for the inferior quadrant.

**Figure 3.**
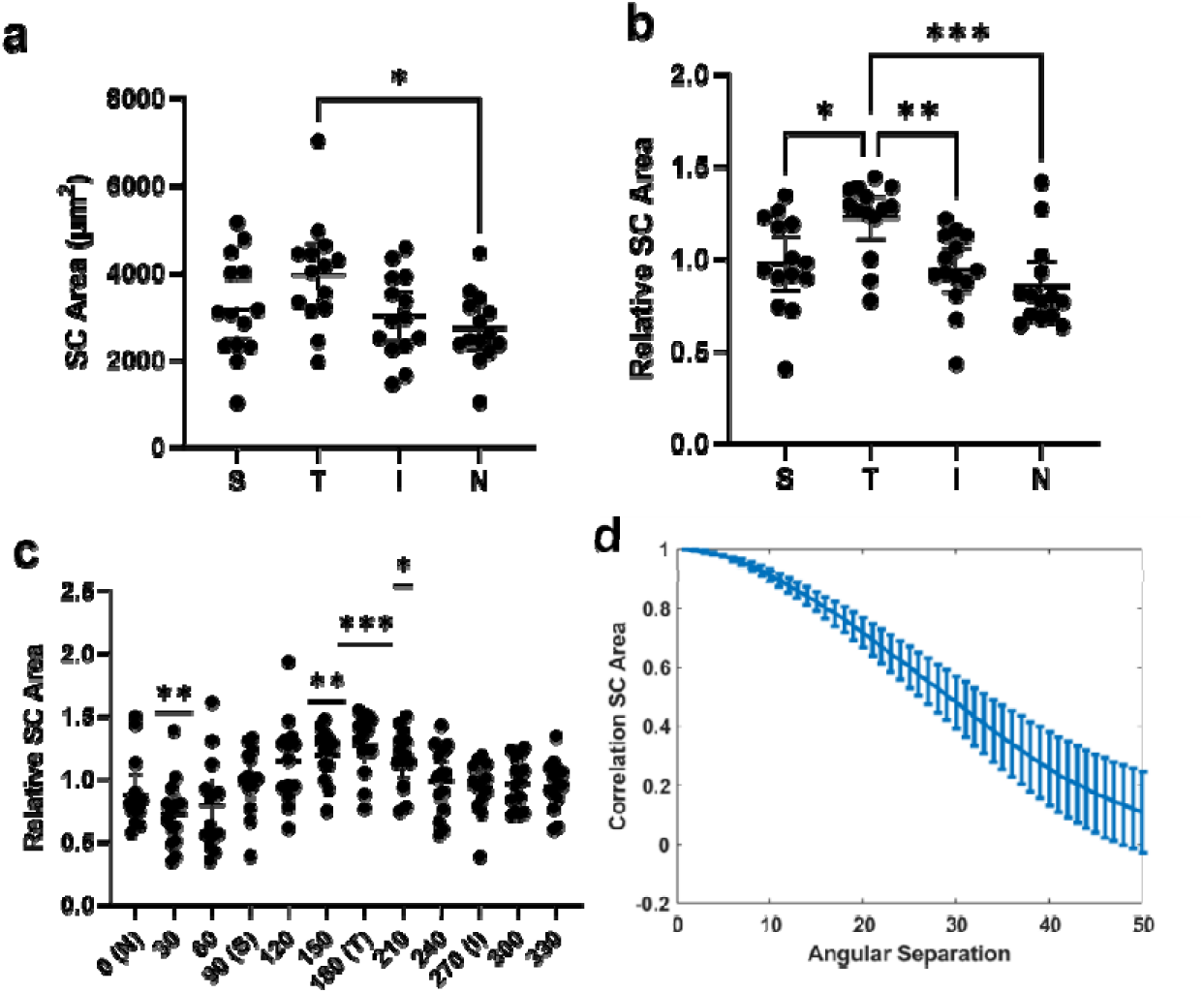
SC has segmental morphology at baseline. **(a)** Average SC area per cross-section for each quadrant. **(b)** Relative SC area for each quadrant normalized to the mean SC area for all quadrants in each eye. **(c)** Relative SC area per 30 degrees normalized to the mean SC area of the eye. Four different sections of 30 degrees had a mean relative SC area statistically different from one. **(d)** Correlation of SC area at a set location with SC area at another location separated by angular separation between 0 and 50 degrees. SC area at cross-sections separated by 50 degrees from each other are uncorrelated. Error bars denote the 95% confidence intervals and n = 14 mice for all baseline analyses. *P < 0.05, ** P < 0.01, ***P < 0.001.

Fig. 3c shows the relative SC cross-sectional area after dividing the eye into twelve finer sub-regions, corresponding to the clock hours of the eye. We found the relative SC cross-sectional area for the eye to be smaller for one clock hour within the nasal quadrant and larger for three clock hours in the temporal quadrant relative to the mean area across all clock hours (p-values: <0.001-0.028). Finally, to assess segmental variation, we tested the correlation between the SC cross-sectional area at a given angular position and the cross-sectional area at adjacent positions. Fig. 3d plots the correlation coefficient between the SC cross-sectional area at two positions separated by angle θ. If SC were homogenous around the limbus, this value would consistently equal 1. Overall, the correlation decreases and drops below 0.2 at 44 degrees.

### Vis-OCT demonstrates change to SC morphology after pilocarpine

We imaged the same eye regions before and after topical pilocarpine or artificial tear administration to assess vis-OCT’s capability to visualize morphological changes in AHO pathways. To assess qualitative changes, we fused B-scans of the same area before and after drug perturbations. Fig. 4a shows an example of a fused image, with B-scans taken at different time points shown in magenta and green, respectively. From the individual images, we generated a composite fused image merging the two individual B-scans. As physical perturbations may alter the outer curvature of the eye, we flattened the fused image such that the outer surface was at the same depth to better visualize differences (Fig. 4b). Fig. 4c shows an example B-scan at baseline (green image) and 15 minutes after pilocarpine administration (magenta image).

**Figure 4.**
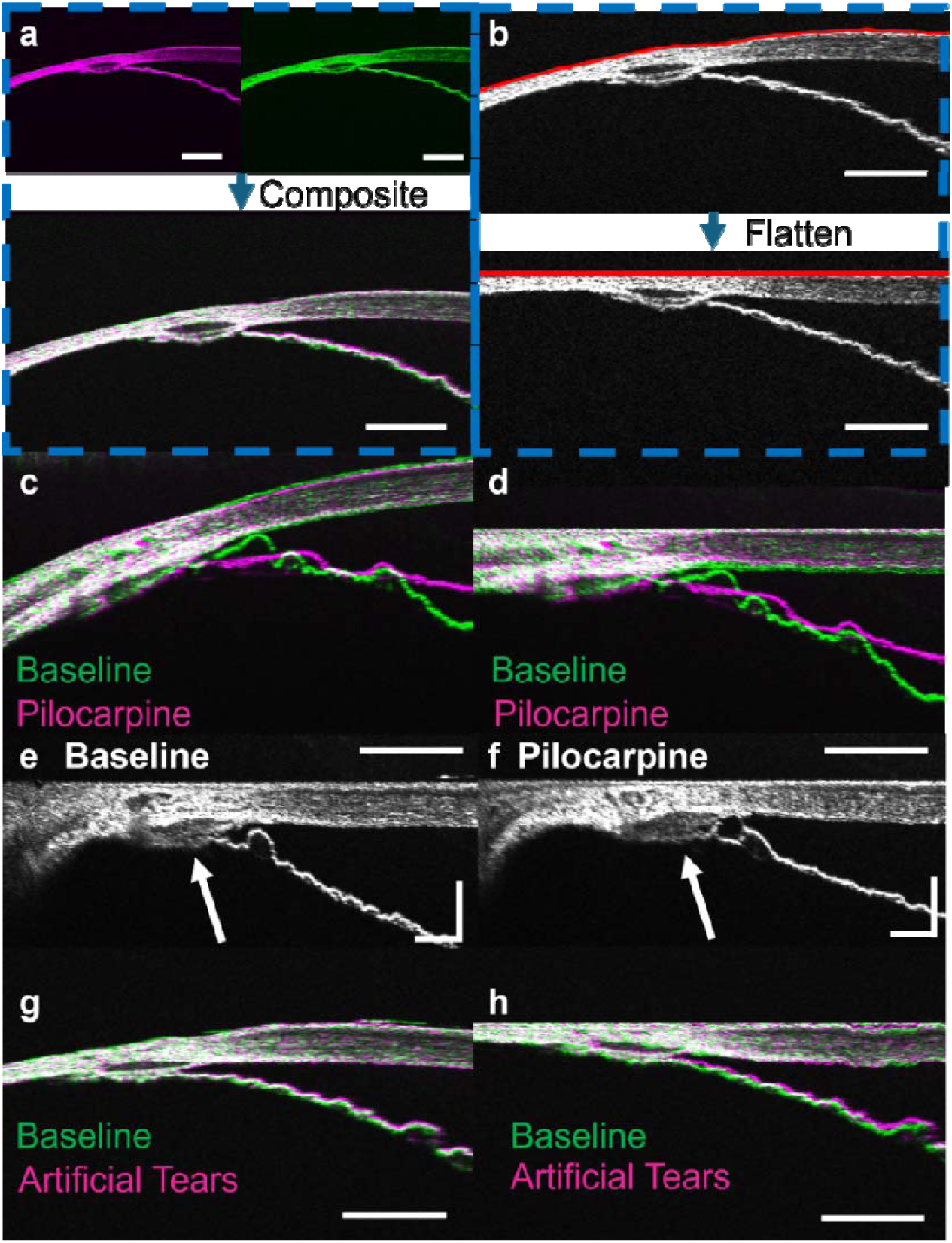
AHO pathway anatomy before and after pilocarpine. **(a)** Composite image of B-scan before and after perturbation generated by overlaying B-scans obtained at different time points (green and magenta images). **(b)** Images are flattened so the eye’s outer surface is at the same depth across the scan. **(c)** Overlaid B-scan at baseline (green) and after pilocarpine administration (magenta). SC size increased after pilocarpine administration, as evidenced by the trabecular borders of SC in the green image (baseline) seen inside the SC lumen of the magenta image (pilocarpine). **(d)** Flattened image shown in (c). **(e)** Another B-scan image before pilocarpine and **(f)** after pilocarpine (arrow: SC). **(g)** B-scan at baseline (green) and after artificial tears (magenta) fused together. No significant changes in SC morphology were observed. **(h)** Flattened image shown in (g). Scale bars are 200 μm, except (e) and (f) which are 100 microns.

We observe that SC is larger after pilocarpine administration, as shown by the outline of the trabecular meshwork (green) for the baseline image being inside that of the pilocarpine image (magenta). If we model SC as an ellipse, we observe that the dimension of the minor and major axes of SC increased in length. Furthermore, we see that natural iris undulations become flatter in response to pilocarpine as would be expected due to pharmacological induction of iris stretching and pupillary miosis (Fig. 4c & d green vs. magenta iris outline). Fig. 4d shows the same B-scans after flattening, with the same qualitative patterns observed as in Fig. 4c. Fig. 4e and f show flattened and uncolored B-scans also demonstrating enlarged SC after pilocarpine (arrows). Fig. 4g shows a B-scan acquired at baseline and after artificial tear administration. We observed no clear differences between the boundaries of SC in this fused image. As seen in the flattened image, we found that baseline and post-eye drop image features aligned closely (Fig. 4h).

Quantitatively, Fig. 5a plots SC volume after pilocarpine and artificial tear administration relative to the volume at baseline. We found that pilocarpine administration increased the SC volume, consistent with our qualitative results and previous studies^43–45^. Relative to the initial volume, pilocarpine increased SC volume (26.8 ± 5.0% increase, p = 0.006, compared to zero), but artificial tears did not (8.9 ± 4.6% increase, p = 0.092, compared to zero). This pilocarpine SC volume increase was greater compared to artificial tears (p = 0.030). We also found that changes in SC volume were more segmental after pilocarpine administration than after artificial tear administration. For each eye, we measured the % volume change for each clock hour and took the standard deviation of the % volume change for each clock hour. Fig. 5b shows the average standard deviation of percent volume change per clock hour, with pilocarpine having a larger change (24.7 ± 3.5%) than after artificial tears (14.3 ± 2.2%) (p = 0.009). Additionally, we found that the % volume change did not impact each quadrant the same. Fig. 5c shows this percent volume change for each quadrant after adding pilocarpine or artificial tears. We saw that the volume increased by 41.1 ± 15.3% in the nasal quadrant, 31.6 ± 8.9% in the superior quadrant, 9.8 ± 5.1% in the temporal quadrant, and 16.4 ± 2.5% in the inferior quadrant after the addition of pilocarpine. The % volume increased by 13.9 ± 4.5% in the nasal quadrant, 1.8 ± 5.7% in the superior quadrant, 13.2 ± 6.7% in the temporal quadrant, and 3.7 ± 5.8% in the inferior quadrant after the addition of artificial tears. The only statistically significant change was seen when pilocarpine was compared to artificial tears for the superior (p = 0.023) and nasal quadrants (p = 0.045).

**Figure 5.**
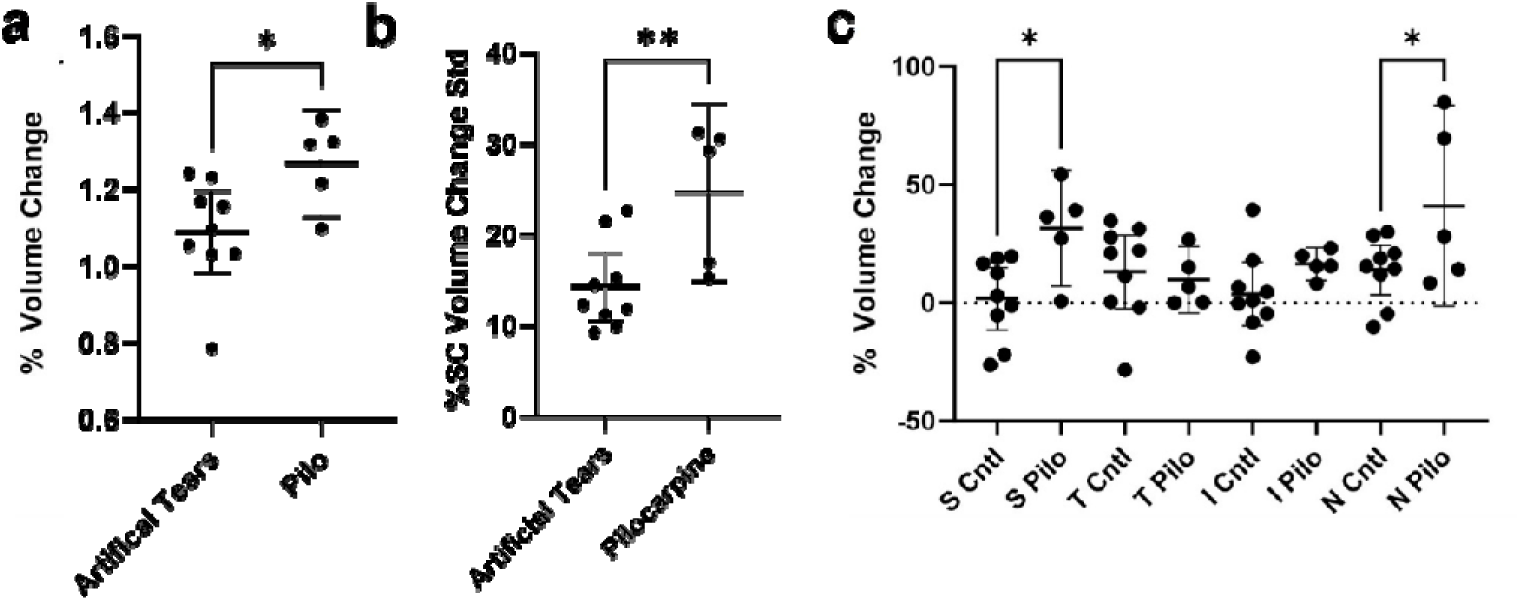
Pilocarpine induced larger SC morphology variation than artificial tears. **(a)** Pilocarpine increased the SC volume relative to artificial tears. **(b)** The average standard deviation in the percent change in SC volume per 30 degrees of the eye was greater in response to pilocarpine than artificial tears. **(c)** The change in percentage SC volume per quadrant was higher in the superior and nasal quadrants when pilocarpine was applied. n = 5 mice for pilocarpine treatment and n = 9 mice for artificial tears application. *P < 0.05, ** P < 0.01.

To further analyze the segmental impact of pilocarpine on SC morphology, we examined the influence of pilocarpine administration on each clock hour of the eye. Fig. 6a shows SC volume per clock hour of the eye normalized to the average volume per clock hour at baseline. SC was larger after pilocarpine addition, with the largest difference in the regions between the nasal and superior quadrants. We saw that the percent volume change was numerically higher for ten out of twelve clock hours after pilocarpine administration than after artificial tear application (Fig. 6b), with the exceptions being two clock hours in the temporal quadrant. We observed that the volume change for pilocarpine was statistically greater for the two clock hours between the nasal and superior quadrant, with volume increases of 52.2 ± 19.6% (p = 0.025) and 55.8 ± 13.3% (p < 0.001), respectively. The corresponding values for artificial tears were 15.2 ± 5.1% and 2.1 ± 5.0%, respectively. Fig. 6c gives the percent SC volume change per clock hour normalized to the average volume across all clock hours. We found that the normalized percent volume change was statistically greater for one clock hour between the nasal and superior quadrants (p = 0.003). Finally, we assessed the correlation between SC volume change at a given angular orientation around the eye and the volume change at a different position separated by an angle θ. Again, if SC were perfectly homogenous around the limbus, this value would consistently equal 1. We found that the correlation decreased and dropped below 0.2 for locations 31 degrees apart.

**Figure 6.**
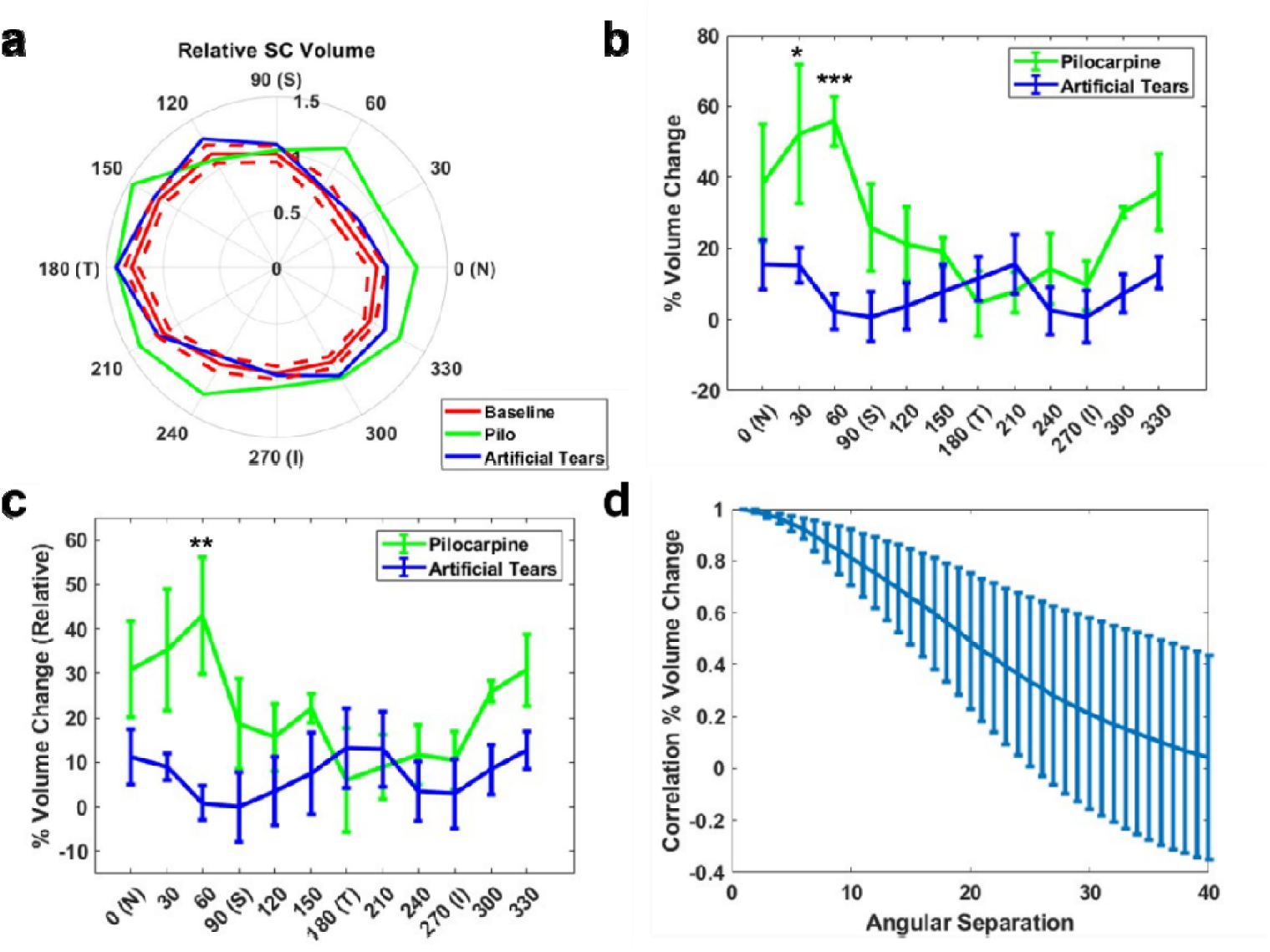
Patterns in SC morphology change per 30 degrees of the eye in response to pilocarpine and artificial tears. **(a)** SC cross-sectional volume per 30 degrees normalized by the mean cross-sectional volume per 30 degrees at baseline for every mouse eye at baseline (red, n = 14) in response to pilocarpine (green, n = 5) and artificial tears (blue, n = 9). The red dotted lines are plus and minus the SEM from the baseline. **(b)** Percentage change in volume for every 30 degrees of the eye. Pilocarpine had the greatest influence in the region between the nasal and superior quadrants. **(c)** Percentage change in volume for every 30 degrees of the eye normalized by the mean volume across all eye regions. **(d)** Correlation of change in SC cross-sectional area at a location with change in cross-sectional area at locations separated by an angular separation between 0 and 40 degrees. Changes in SC area between locations separated by greater than 30 degrees are uncorrelated. All error bars are SEM. *P < 0.05, ** P < 0.01, ***P < 0.001.

## Discussion

In this work, we evaluated circumferential AHO anatomy and physiological drug response *in vivo* for the first time. Using a robotic vis-OCT system, a full 3D and circumferential assessment of mouse SC before and after topical pilocarpine administration was made. Segmental SC anatomy was observed at baseline in different mice, and on average, SC was larger in the temporal quadrant while smaller in the nasal quadrant (Fig. 3). After pilocarpine administration, SC also became larger in size (Fig. 4), and this change was segmental as it was more pronounced in the superior and nasal quadrants (Fig. 5).

Pilocarpine has been used for decades in glaucoma patients to lower IOP^38, 39, 46^. Pilocarpine is a muscarinic receptor agonist that increases AHO facility through the trabecular pathways. Pilocarpine constricts the longitudinal ciliary muscle to pull down on the scleral spur and widen the TM^38^. Multiple studies have shown in live humans and live mice that pilocarpine administration leads to a larger SC^43, 44^. However, in these studies, the imaging was restricted to certain regions of the eye, and a full 360-degree circumferential assessment has never been performed. The unique finding in this study was not just that baseline SC was circumferentially segmental but that the SC response to pilocarpine was segmental as well.

The segmental SC response to pilocarpine and baseline SC segmental appearance raises important considerations. First, this result is consistent with preliminary aqueous angiography imaging in humans using Miochol-E. Miochol-E is acetylcholine (ACh) that is FDA-approved for intracameral application, and patients are known to have lower post-operative IOP after Miochol-E use during cataract surgery^47^. This makes sense as ACh is the endogenous muscarinic neurotransmitter (which pilocarpine mimics) that contracts the ciliary muscle. Early results using aqueous angiography have shown that Miochol-E leads to segmental improvement of angiographic AHO in focal areas (Huang AS., et al., 2022, Abstract, American Glaucoma Society conference). Combined with the current mouse research results, one question is why a trabecular meshwork drug response can be segmental. As small molecules, pilocarpine and ACh should diffuse throughout the eye and impact the TM circumferentially and uniformly. However, segmental LF TM regions have shown differential gene/protein expression^9, 11, 12^, ECM deposition^10, 11^, and increased biomechanical stiffness^12^. This may explain the variable circumferential response to the drug. For example, LF TM may be less amenable to longitudinal ciliary muscle contraction initiated by muscarinic agonists if there is increased pro-fibrotic protein expression, ECM deposition, and tissue stiffness. Further, it may be that the expansion of baseline LF AHO regions and LF characteristics lead to ocular hypertension (OHTN) in glaucoma. Then, differing degrees of these LF characteristics in any particular patient or eye may explain variable responses to IOP-lowering medications.

The full circumferential imaging of AHO pathways in this work is also critical for studying segmental AHO anatomy and drug response. Using robotic AS-OCT, the circumferential reconstruction of AHO pathways effectively created a “digital twin” of the anterior segment and AHO pathways. This approach and the “digital twin” hold promise to improve the understanding of AHO physiology/pathophysiology and to improve glaucoma clinical care. Aqueous angiography was developed for circumferential AHO imaging in patients. For glaucoma patients and surgery, aqueous angiography-targeted trabecular surgery to baseline LF regions has shown AHO rescue^27^. In an anterior segment perfusion system, aqueous angiography-targeted trabecular surgery to baseline LF regions also resulted in greater AHO facility increase and IOP reduction compared to surgery in baseline HF regions^21^. Future clinical studies are planned to confirm this. However, aqueous angiography is expensive, laborious, time-consuming, and invasive. With a 360-degree “digital twin” of AHO pathways, non-invasive pre- or intra-operative AS-OCT imaging could be performed instead to guide glaucoma surgery. However, the missing knowledge gap is a structure/function relationship and the identification of OCT- imaged structural biomarkers that predict angiographic LF and HF regions. Thus, in the future, the development of a clinical robotic AS-OCT dedicated to AHO assessment is planned to create eye-specific anterior segment “digital twins” in humans that can be compared to gold-standard aqueous angiography imaging to find structural proxies for segmental AHO.

There are several limitations in this study. First, the sample size is limited. Also, IOP was not measured. However, the mouse AHO facility and IOP response to pilocarpine are well-studied^43, 48^, and our results demonstrated a flattening of the iris contour consistent with an expected pilocarpine miotic pupillary response (Fig. 4). Also, species-specific and technical factors must be considered before direct translation of this work to humans. In humans, AHO is known to be the greatest nasal^26, 49^ which was not seen here (Fig. 3). Mouse imaging required systemic anesthetics and relaxing incisions on the eyelids to visualize SC on a posteriorly positioned mouse limbus, which could have impacted IOP and AHO. For humans, the limbus is more anterior on the eye and thus more easily visualized in awake, unperturbed, and relaxed forward-looking individuals. Thus, if eyes are widely opened, clinical robotic AS-OCT imaging of AHO pathways may not have the same challenges as in mouse eyes.

In conclusion, high-resolution circumferential imaging of the anterior segment and AHO pathways is possible using robotic AS-OCT. Observation of segmental SC anatomy is consistent with known segmental AHO seen in flow-based imaging studies. Segmental AHO anatomical response to a muscarinic agonist was seen, and this result opens the door to a better understanding of baseline segmental AHO pathway characteristics. Studying drug-responsive and non-responsive regions may lead to an improved understanding of how OHTN arises in the first place, why there is variable patient response to IOP-lowering drugs, and identification of potential new drug targets by specifically understanding “rescuable” regions. Circumferential imaging may also allow for OCT-only determination of segmental LF and HF regions in the future. With this knowledge, there is the potential for not only improved glaucoma pharmacological and surgical treatment but personalized therapy.

## Acknowledgments

Funding for this work came from National Institutes of Health, Bethesda, MD (Grant Numbers U01EY033001 [HFZ], R01EY033813 [HFZ], R01EY034740 [HFZ], R01EY030501 [ASH], P30EY022589 UCSD core grant, F30EY034033 [RF]), Research to Prevent Blindness David Epstein Career Advancement Award in Glaucoma Research sponsored by Alcon [ASH] and an unrestricted grant from Research to Prevent Blindness (New York, NY) [UCSD].

## Disclosures

Abbvie/Allergan (AH; C), Amydis (AH; C), Celanese (AH; C), Diagnosys (AH; research support), Equinox (AH; C), Glaukos (AH; C & R), Heidelberg Engineering (AH; R), QLARIS (AH; C), Santen (AH; C), and Topcon (AH; C). Opticent (HFZ; O&P; CS; O&P)

## References

1. Weinreb RN, Khaw PT. Primary open-angle glaucoma. Lancet 2004;363:1711–1720.

2. Kass MA, Heuer DK, Higginbotham EJ, et al. The Ocular Hypertension Treatment Study: a randomized trial determines that topical ocular hypotensive medication delays or prevents the onset of primary open-angle glaucoma. Arch Ophthalmol 2002;120:701–713; discussion 829-730.

3. Musch DC, Gillespie BW, Niziol LM, Lichter PR, Varma R, Group CS. Intraocular pressure control and long-term visual field loss in the Collaborative Initial Glaucoma Treatment Study. Ophthalmology 2011;118:1766–1773.

4. The Advanced Glaucoma Intervention Study (AGIS): 7. The relationship between control of intraocular pressure and visual field deterioration.The AGIS Investigators. Am J Ophthalmol 2000;130:429-440.

5. Brubaker RF. Goldmann’s equation and clinical measures of aqueous dynamics. Exp Eye Res 2004;78:633–637.

6. Grant WM. Experimental aqueous perfusion in enucleated human eyes. Arch Ophthalmol 1963;69:783–801.

7. Johnson M. What controls aqueous humour outflow resistance. Experimental Eye Research 2006;82:545–547.

8. M J, K E. Mechanisms and routes of aqueous humor drainage. In: DM A, FA J (eds), Principles and practice of ophthalmology. Philadelphia: WB Saunders; 2000.

9. Saraswathy S, Bogarin T, Barron E, et al. Segmental differences found in aqueous angiographic-determined high - and low-flow regions of human trabecular meshwork. Exp Eye Res 2020;196:108064.

10. Keller KE, Bradley JM, Vranka JA, Acott TS. Segmental versican expression in the trabecular meshwork and involvement in outflow facility. Invest Ophthalmol Vis Sci 2011;52:5049–5057.

11. Vranka JA, Bradley JM, Yang YF, Keller KE, Acott TS. Mapping molecular differences and extracellular matrix gene expression in segmental outflow pathways of the human ocular trabecular meshwork. PLoS One 2015;10:e0122483.

12. Vranka JA, Staverosky JA, Reddy AP, et al. Biomechanical Rigidity and Quantitative Proteomics Analysis of Segmental Regions of the Trabecular Meshwork at Physiologic and Elevated Pressures. Invest Ophthalmol Vis Sci 2018;59:246–259.

13. Battista SA, Lu Z, Hofmann S, Freddo T, Overby DR, Gong H. Reduction of the available area for aqueous humor outflow and increase in meshwork herniations into collector channels following acute IOP elevation in bovine eyes. Invest Ophthalmol Vis Sci 2008;49:5346–5352.

14. Saraswathy S, Tan JC, Yu F, et al. Aqueous Angiography: Real-Time and Physiologic Aqueous Humor Outflow Imaging. PLoS One 2016;11:e0147176.

15. Huang AS, Saraswathy S, Dastiridou A, et al. Aqueous Angiography with Fluorescein and Indocyanine Green in Bovine Eyes. Transl Vis Sci Technol 2016;5:5.

16. Huang AS, Saraswathy S, Dastiridou A, et al. Aqueous Angiography-Mediated Guidance of Trabecular Bypass Improves Angiographic Outflow in Human Enucleated Eyes. Invest Ophthalmol Vis Sci 2016;57:4558–4565.

17. Snyder KC, Oikawa K, Williams J, et al. Imaging Distal Aqueous Outflow Pathways in a Spontaneous Model of Congenital Glaucoma. Transl Vis Sci Technol 2019;8:22.

18. Burn JB, Huang AS, Weber AJ, Komáromy AM, Pirie CG. Aqueous Angiography in Normal Canine Eyes. Transl Vis Sci Technol 2020;9:44.

19. Burn JB, Huang AS, Weber A, Komáromy AM, Pirie CG. Aqueous angiography in pre-glaucomatous and glaucomatous ADAMTS10-mutant canine eyes: A pilot study. Vet Ophthalmol 2022;25 Suppl 1:72–83.

20. Huang AS, Li M, Yang D, Wang H, Wang N, Weinreb RN. Aqueous Angiography in Living Nonhuman Primates Shows Segmental, Pulsatile, and Dynamic Angiographic Aqueous Humor Outflow. Ophthalmology 2017.

21. Strohmaier CA, Wanderer D, Zhang X, et al. Greater Outflow Facility Increase After Targeted Trabecular Bypass in Angiographically Determined Low-Flow Regions. Ophthalmol Glaucoma 2023.

22. Strohmaier CA, McDonnell FS, Zhang X, et al. Differences in Outflow Facility Between Angiographically Identified High- Versus Low-Flow Regions of the Conventional Outflow Pathways in Porcine Eyes. Invest Ophthalmol Vis Sci 2023;64:29.

23. Dada T, Bukke AN, Huang AS, Sharma N, Verma S. Recruitment of Temporal Aqueous Outflow Channels After Bent Needle Ab-Interno Goniectomy Demonstrated by Aqueous Angiography. J Glaucoma 2023;32:e15–e18.

24. Gupta S, Zhang X, Panigrahi A, et al. Reduced Aqueous Humor Outflow Pathway Arborization in Childhood Glaucoma Eyes. Transl Vis Sci Technol 2024;13:23.

25. Huang AS, Penteado RC, Saha SK, et al. Fluorescein Aqueous Angiography in Live Normal Human Eyes. J Glaucoma 2018.

26. Huang AS, Camp A, Xu BY, Penteado RC, Weinreb RN. Aqueous Angiography: Aqueous Humor Outflow Imaging in Live Human Subjects. Ophthalmology 2017.

27. Huang AS, Penteado RC, Papoyan V, Voskanyan L, Weinreb RN. Aqueous Angiographic Outflow Improvement after Trabecular Microbypass in Glaucoma Patients. Ophthalmol Glaucoma 2019;2:11–21.

28. Rosenfeld PJ, Windsor MA, Feuer WJ, et al. Estimating Medicare and Patient Savings From the Use of Bevacizumab for the Treatment of Exudative Age-related Macular Degeneration. Am J Ophthalmol 2018;191:135–139.

29. CMS. Top 200 Level 1 Cpt Codes Ranked by Charges. 2019.

30. Hann CR, Fautsch MP. Preferential fluid flow in the human trabecular meshwork near collector channels. Invest Ophthalmol Vis Sci 2009;50:1692–1697.

31. Hann CR, Bentley MD, Vercnocke A, Ritman EL, Fautsch MP. Imaging the aqueous humor outflow pathway in human eyes by three-dimensional micro- computed tomography (3D micro-CT). Exp Eye Res 2011;92:104–111.

32. Huang AS, Belghith A, Dastiridou A, Chopra V, Zangwill LM, Weinreb RN. Automated circumferential construction of first-order aqueous humor outflow pathways using spectral-domain optical coherence tomography. J Biomed Opt 2017;22:66010.

33. Kagemann L, Wollstein G, Ishikawa H, et al. 3D visualization of aqueous humor outflow structures in-situ in humans. Exp Eye Res 2011;93:308–315.

34. Skaat A, Rosman MS, Chien JL, et al. Effect of Pilocarpine Hydrochloride on the Schlemm Canal in Healthy Eyes and Eyes With Open-Angle Glaucoma. JAMA Ophthalmol 2016;134:976–981.

35. Skaat A, Rosman MS, Chien JL, et al. Microarchitecture of Schlemm Canal Before and After Selective Laser Trabeculoplasty in Enhanced Depth Imaging Optical Coherence Tomography. J Glaucoma 2017;26:361–366.

36. Wang F, Shi G, Li X, et al. Comparison of Schlemm’s canal’s biological parameters in primary open-angle glaucoma and normal human eyes with swept source optical. J Biomed Opt 2012;17:116008.

37. Fang R, Zhang P, Zhang T, et al. Freeform robotic optical coherence tomography beyond the optical field-of-view limit. *bioRxiv* 2024.

38. Barany EH. The mode of action of pilocarpine on outflow resistance in the eye of a primate (Cercopithecus ethiops). Invest Ophthalmol 1962;1:712–727.

39. Bartels SP, Neufeld AH. Mechanisms of topical drugs used in the control of open angle glaucoma. Int Ophthalmol Clin 1980;20:105–116.

40. Fang R, Zhang P, Zhang T, et al. Freeform robotic optical coherence tomography beyond the optical field-of-view limit. *bioRxiv* 2024;2024.2005.2021.595073.

41. Zhang X, Beckmann L, Miller DA, et al. In Vivo Imaging of Schlemm’s Canal and Limbal Vascular Network in Mouse Using Visible-Light OCT. Invest Ophthalmol Vis Sci 2020;61:23.

42. Zhang P, Miller EB, Manna SK, Meleppat RK, Pugh EN, Jr., Zawadzki RJ. Temporal speckle-averaging of optical coherence tomography volumes for in-vivo cellular resolution neuronal and vascular retinal imaging. Neurophotonics 2019;6:041105.

43. Li G, Farsiu S, Chiu SJ, et al. Pilocarpine-induced dilation of Schlemm’s canal and prevention of lumen collapse at elevated intraocular pressures in living mice visualized by OCT. Invest Ophthalmol Vis Sci 2014;55:3737–3746.

44. Skaat A, Rosman MS, Chien JL, et al. Effect of Pilocarpine Hydrochloride on the Schlemm Canal in Healthy Eyes and Eyes With Open-Angle Glaucoma. JAMA Ophthalmology 2016;134:976–981.

45. Kinney ML, Johnson AD, Reddix M, McCann MB. Temporal Effects of 2% Pilocarpine Ophthalmic Solution on Human Pupil Size and Accommodation. Military Medicine 2020;185:435–442.

46. Drance SM, Nash PA. The dose response of human intraocular pressure to pilocarpine. Can J Ophthalmol 1971;6:9–13.

47. West J, Burke J, Cunliffe I, Longstaff S. Prevention of acute post-operative pressure rises in glaucoma patients undergoing cataract extraction with posterior chamber lens implant. Br J Ophthalmol 1992;76:534–537.

48. Overby DR, Bertrand J, Schicht M, Paulsen F, Stamer WD, Lütjen-Drecoll E. The structure of the trabecular meshwork, its connections to the ciliary muscle, and the effect of pilocarpine on outflow facility in mice. Invest Ophthalmol Vis Sci 2014;55:3727–3736.

49. Johnstone M, Jamil A, Martin E. Aqueous Veins and Open Angle Glaucoma. In: Schacknow PN, Samples JR (eds), The Glaucoma Book: A Practical, Evidence- Based Approach to Patient Care. New York: Springer; 2010:65-78.

